# Followers of honeybee waggle dancers change their behavior when dancers are sleep-restricted or perform imprecise dances

**DOI:** 10.1101/457598

**Authors:** Barrett A. Klein, Michael Vogt, Keaton Unrein, David M. Reineke

## Abstract

Sleep loss is known to adversely affect language use in humans (Shein, 1957; Morris, Williams, & Lubin, 1960; Horne, 1988; Wimmer, Hoffmann, Bonato, & Moffitt, 1992; Harrison & Horne, 1997; Harrison & Horne, 1998), but evidence for effects on language comprehension is sparse (Schein, 1957; Pilcher et al., 2007; Pilcher, Jennings, Phillips, & McCubbin, 2016) and we are aware of no studies explicitly examining the effects of sleep loss on communication – both signaling and receiving – in any other species. This includes studies of signals affected by sleep loss and the behavioural consequences exhibited by receivers of these signals. Possible exceptions relate to courtship (Seugnet et al., 2011) and aggression (Kayser, Mainwaring, Yue, & Sehgal, 2015), although studies will typically give little or no detail about the behaviour of receivers. We know almost nothing about the ecology of sleep in invertebrates (Tougeron & Abram, 2017), or the consequences of sleep loss on invertebrate communication, hampering our understanding of basic operations performed by the majority of animal species.

Communication is effective only when a reliable signal is delivered and properly received. The perfect signal can fall on deaf ears, or an imperfect signal can confuse or mislead an intended recipient. Unavoidable errors exist in communication, and can result from unreliable signaling, even in the absence of deception (Carazo & Font, 2014). How differences in signal precision arise and how potential receivers respond to error in a signal are still largely unknown, although responses to impaired signaling can reveal fitness-relevant costs, be it in the competitive signal disruption of treehoppers during courtship (Legendre, Marting, & Cocroft, 2012), response to predator or prey stimuli by social spiders (Pruitt et al., 2016), or, in studies designed to impair communication, the attempted recruitment of honeybees by dancers (Dornhaus, 2002; Sherman & Visscher, 2002).

Communication is extensively documented in western honeybees (*Apis mellifera*), thanks to Karl von Frisch’s seminal work decoding the waggle dance (von Frisch, 1967). A waggle dance performed by *A. mellifera* foragers consists of a sequence of circuits on a vertical comb, capable of advertising the direction, distance, and quality of an advertised site. Returning from a foraging trip kilometers from her nest, a worker bee can communicate to her nest mates the location of a food source by performing a body-waggling motion at an angle relative to the vertical (waggle phase), which corresponds to the angle of the food source relative to the sun’s azimuth. Iterations of these stereotyped motions on the comb, consisting of alternating waggle phases and return phases, result in error, and angular precision varies across dances (De Marco, Gurevitz, & Menzel, 2008). If a small insect advertises a food source 1 km away, it would seem important to deliver an accurate message with a high degree of precision.

Understanding the relationship between communication and sleep depends on the use of a clear, consistent description of both communication and sleep. Definitions of communication vary, particularly in terms of intentionality and information (Rendall, Owren, & Ryan, 2009). A signaler may intend on influencing a receiver, or consequences of signaling may be inadvertent (Seyfarth & Cheny, 2003). For our purposes, communication is simply a social event that includes a signaler (forager honeybee performing waggle dances) and a receiver (dance follower) behaviourally responding to a signal.

Sleep is relatively well documented in honeybees, with studies identifying sleep dynamics (Sauer, Kinkelin, Herrmann, & Kaiser, 2003), plasticity (Klein & Seeley, 2011), and caste-dependent behaviour (Eban-Rothschild & Bloch, 2008; Klein, Olzsowy, Klein, Saunders, & Seeley, 2008), and can be identified by a suite of behaviours, including increased response threshold to external stimuli (Kaiser, 1988), and internal control, with restriction of the condition resulting in a rebound of the behaviour (Sauer, Herrmann, & Kaiser, 2004). Klein et al. (2010) used superficial indicators of sleep in honeybees [(relative immobility and discontinuous ventilation of the metasoma; see Klein et al. (2008)] to show that sleep-restricted foragers dance with less precision in the direction component of the dance (measured as error around the mean angle of a dance) relative to control bees. If sleep loss (potentially caused by the actions of predators, parasites, abiotic factors, or beekeeping practices) degrades signaling, an obvious consequence could involve decreased colony fitness. No information, however, was gathered by Klein et al. (2010) concerning the actions of the receivers (dance followers). Knowing how receivers respond to imprecise signals is a critical step to understanding colony-level fitness consequences of imprecision in this model system. If different honeybees perform dances for the same distant food source, how do followers of relatively imprecise dances respond? We analysed the actions of dance followers to see if following a relatively imprecise dance results in switching dancers rather than exiting the nest, a response we predicted when receivers are confronted with a potentially costly signal error. We also looked at sleep restriction of dancers as a factor affecting dance follower behaviour.

## Methods

### Study site and experimental design

Klein et al. (2010) outfitted an observation hive of Italian honeybees (*Apis mellifera ligustica* Spinola, 1806) with a device called the insominator to induce sleep loss in a select subset of dancers in the nest, and video-recorded the effects of sleep loss on these dancers relative to control dancers. In this study, we use the same video recordings to investigate activities of dance followers. Foragers were trained to visit a feeder offering 1.0 M sucrose solution scented with anise each day of the study, 1 km from the nest at Cranberry Lake Biological Station in Adirondack State Park, NY (44°09’N, 74°48’W). We selected this site because the surrounding forest offers few natural food sources for honeybees, making our feeder attractive, and to foragers exclusively from our colony (Wray et al., 2008). We marked a forager’s mesothorax with either ferrous steel (Fe) tags or nonferrous copper (Cu) tags. The manually-operated insominator slid along a track, with a bank of magnets on each side of the observation hive, jostling only bees with ferrous steel tags. The insominator did not noticeably disturb copper-tagged control bees. Tagged bees’ waggle dances were video recorded (Panasonic AGDVC 30, Kadoma, Japan) and these dances were analysed for effects of directional precision, before and after restricting sleep of treatment bees. We used videos taken during the day immediately following a 12-h nocturnal period of insominator operation to analyse unmarked members of the colony that followed dances performed by the tagged dancers.

### Dancers and dance followers

Videos of the “dance floor” captured almost all dances by each forager, including the metal-tagged foragers trained to our feeder. A tunnel, located immediately off-screen in the lower right corner of the video frame, served as the only nest entrance, allowing us to record when bees likely exited the nest. Individually-marked, metal-tagged foragers performed waggle dances, with each dance consisting of a series of waggle phases. Directional precision of each dance was measured as the standard deviation (SD), in degrees, around the mean across all waggle phases performed in a dance. We selected a subset of relatively precise dances (SD <10.45°) and imprecise dances (SD ≥16.45°) to facilitate our analysis of follower behaviour based on precision of dances followed (Fig 1). Gathering data from every follower of every dance recorded was temporally prohibitive, so by excluding dances in between these two subsets, we aimed to more economically assess effect of high precision versus low precision on follower behaviour. Our selection of dances was also an effort to maximize the sample size of unique dancers. When selecting dances, we attempted to control for number of waggle phases, but because our sample did not allow for this, we also tested for the effect of the number of waggle phases per dance in follower behaviour.

**Fig 1.**
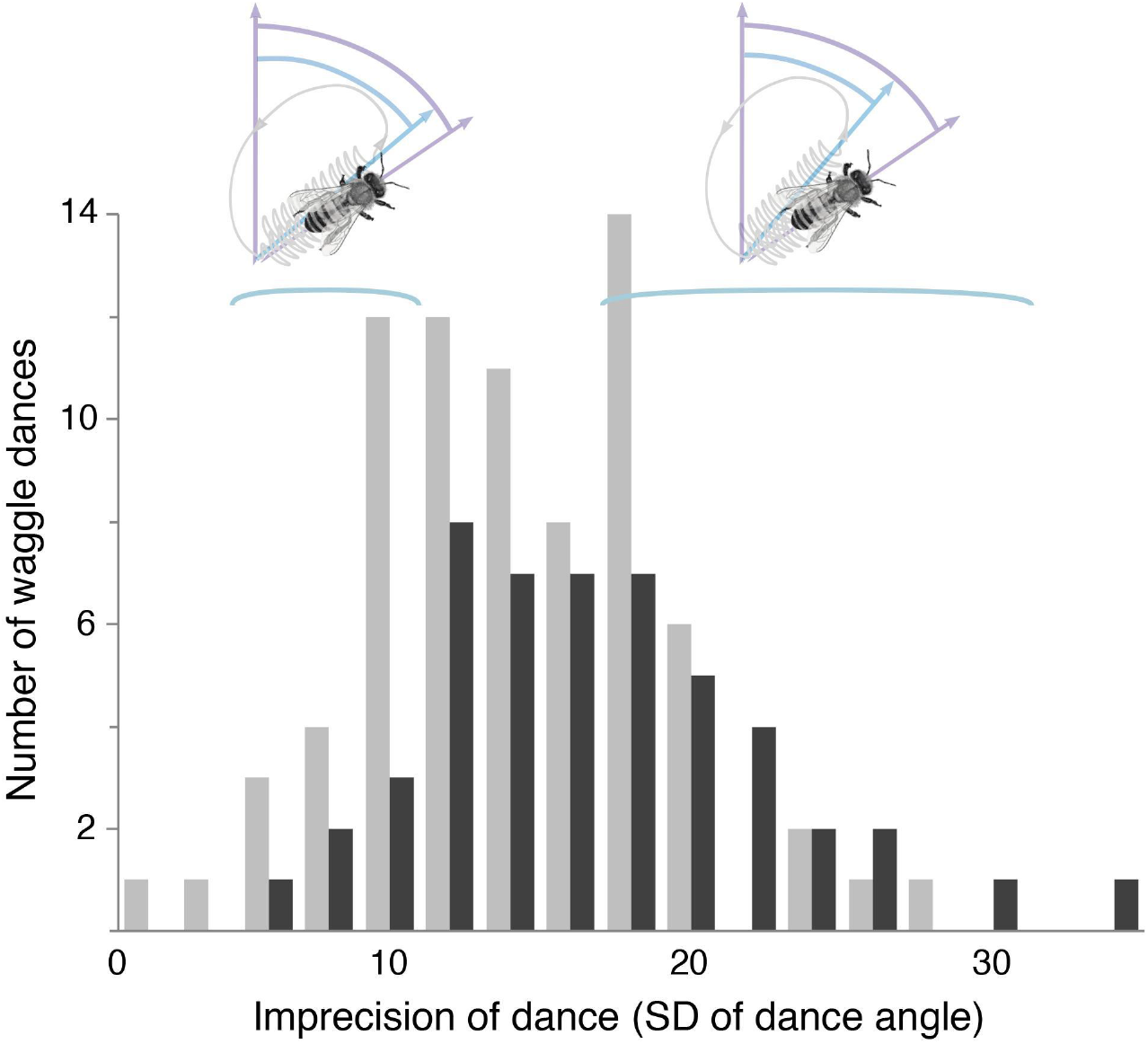
Frequency distribution of dances with high to low precision, as measured by the standard deviations (SD) around the mean of dance angles (representing precision of direction component of waggle dance). Dances were performed by steel-tagged bees (black bars) and copper-tagged bees (gray bars) (*N* = 126 dances), following sleep restriction of steel-tagged bees (Klein et al., 2010). Highlighted areas represent bins from which dances were selected for analysis because these dances were relatively precise (left, with less variance in dance angle, pictured above) or imprecise (right, with greater variance in dance angle, pictured above).

We were initially interested in how followers responded to dance precision more than how the followers responded to sleep-restricted dancers, so we first compared precision between sleep-restricted treatment bees and control bees within precise and imprecise sets of dances, above, to see if we could combine data across bees within each group (precision bin) of dances. Sleep restriction resulted in less precise dances overall (Klein et al., 2010), but as we predicted, the average measure of precision did not differ between treatment and control bees within the precise bin (8.1 ± 1.0° vs. 8.2 ± 0.5°, *N* = 6 & 21 dances, respectively; *z* = −0.06, *P* = 0.998) or within the imprecise bin (21.3 ± 0.9° vs. 19.3 ± 0.9°, *N* = 22 & 24 dances, respectively; *z* = 1.59, *P* = 0.18), so data from sleep-restricted and control bees were combined within bins of dances (Fig 1). Dance data were analysed with precision as continuous measure, except where noted.

For the purpose of studying dance follower behaviour, we examined waggle dances consisting of minimally three waggle phases. We identified followers of dances by adopting criteria used by Wray et al. (2008): The potential follower must (1) be within one bee’s body length of the dancer (Judd, 1995), (2) be facing in the general direction of the dancer (whether facing behind or to the side of a dancer appears to be irrelevant; Tanner & Visscher, 2009; Gil & De Marco, 2010), and (3) follow at least two complete waggle phases, so as to distinguish from a bee simply walking past the dancer (Wray et al., 2008), or a nectar receiver uninterested in the location advertised (Toufailia, Couvillon, Ratnieks, & Grüter, 2013) (Fig 2). Each follower of a dance was unique, and even though all unmarked bees arriving at the feeder were captured, we cannot be certain that all follower events across dances were by unique bees.

**Fig 2.**
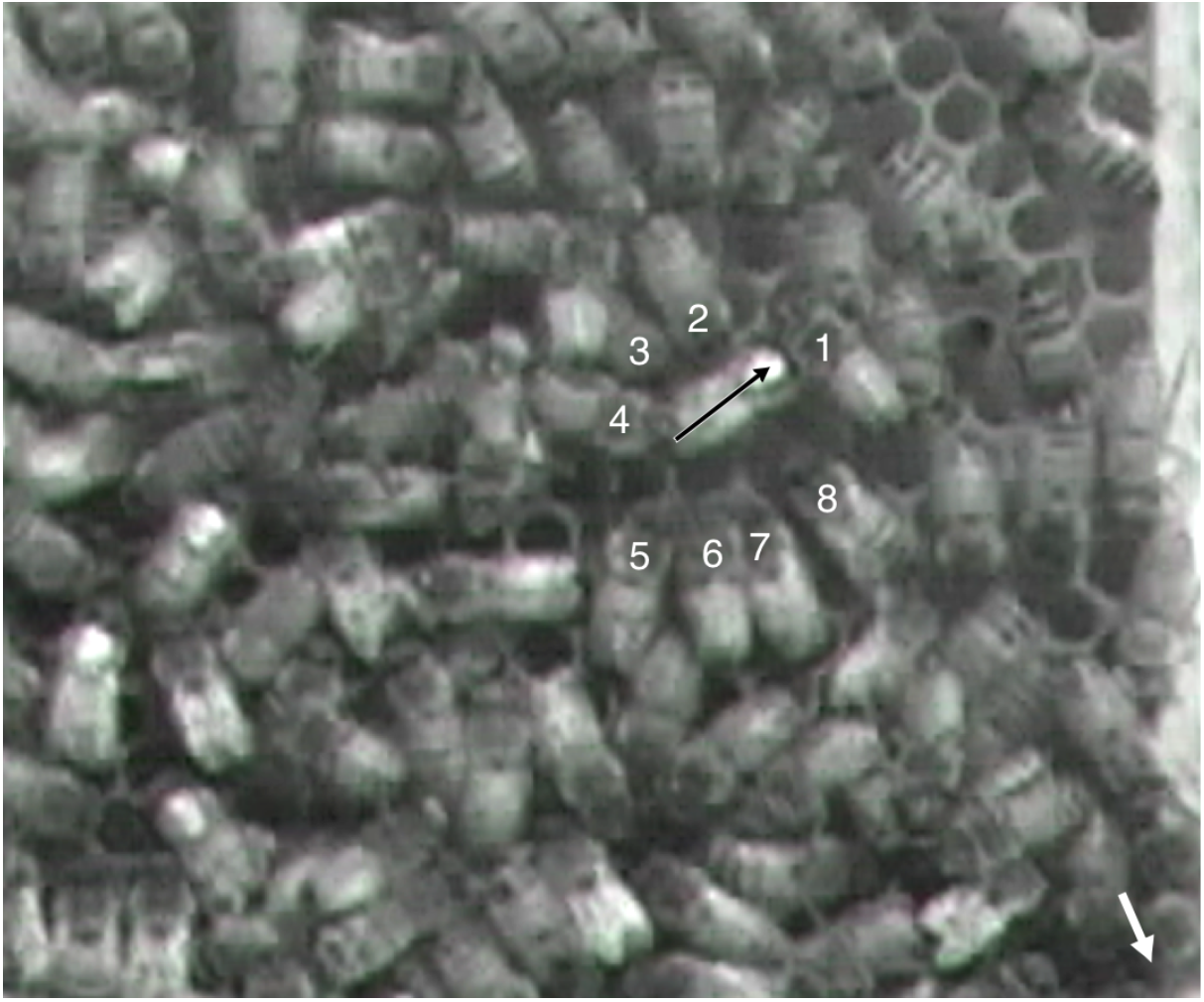
Detail from video still, with eight worker honeybees following an individually paint-marked dancer. Direction of dance is indicated by black arrow, aligned with dancer’s body axis; the black arrowhead points to painted metal tag, which was adhered to the center of the dancer’s dorsal mesothorax. Followers (numbers 1-8) were designated as being within one bee’s body length of dancer, oriented toward dancer, and having followed at least two waggle phases of the dance. Note three other individually-marked foragers with light-colored tags at lower left. Entrance of nest is located below lower right corner of image, indicated by the white arrow. Behaviour of every dance follower of pre-selected dances was recorded.

### Data collection and analysis

Marked dancers attracted multiple unmarked followers (Fig 2). Using VirtualDub (v.1.10.4), we modified playback speeds and recorded screenshots each time a dancer performed a waggle phase to identify and keep track of dance followers. Authors MV, KU, and helpers JK and SS collected data blind to a dancer’s identity, and inter-observer error for calculating number of waggle phases followed was 0.91 ± 0.46 phases. Data collected include unique identity of dancer followed, number of waggle phases per dance followed by each dance follower, duration of each following event (ultimately not used because added no new information; data correlated strongly with number of waggle phases followed), and whether the follower exited the nest (Movie S1, clip 1), switched to another dancer (for a minimum of two complete waggle phases; Movie S1, clip 2), or we lost her location. Data related to signalers (dancers) were made available by Klein et al. (Klein et al., 2010). Results include means ± SE, unless otherwise noted. We set α = 0.05 for all tests, and report *P*-values as two-tailed. DMR performed all statistical tests using R (R Foundation for Statistical Computing, 2015) and lme4 (Bates, Mächler, Bolker, & Walker, 2015) with lsmeans (Lenth, 2016) to perform a linear mixed effects logistic regression analysis of the relationship between SD of the angle and the fixed effect treatment, and for the switching vs. exiting analysis. As a random effect, we allowed for separate intercepts for dancer bees. Visual inspection of residual plots did not reveal any obvious deviations from homoscedasticity or normality for any of the test results that rely on these conditions. The *P*-value was obtained by a general linear hypothesis test. We treated multiple dances performed by the same dancer as subsamples of that dancer (experimental unit) to avoid pseudoreplication. From the perspective of a dance follower, however, each dance represented a discrete and unique set of information, so the dance, rather than the dancer, was treated as the experimental unit. Thus, for analyses of dance followers (e.g., number of waggle phases followed), we treated followers of different dances as independent, whereas we treated followers of the same dance as subsamples of that dance (Datasets S1-S8, Code S1).

## Results

We analysed the behaviour of 615 follower events of 39 waggle dances performed by 13 dancers (Table 1).

**Table 1.** Dance and dance follower data, binned by relative precision (precise: SD <10.45°, imprecise ≥16.45°) of dance x dancer group (Cu-tagged dancers, and sleep-restricted, Fe-tagged dancers). We selected a subset of dances with followers, and recorded behaviour during and after a following event, although some bees were “lost” due to tracking constraints. Mean numbers include ± SE.

| Dances | # waggle phases<br>(n = # dances) | # followers<br>(n = # dances with followers) | # waggle phases followed<br>(n = # followers) | % switch<br>(n = # followers, lost excluded) | % exit<br>(n = # followers, lost excluded) | # dances | # dances with followers | # followers | # followers, lost excluded |
| --- | --- | --- | --- | --- | --- | --- | --- | --- | --- |
| Cu-Precise | 11.3 ± 2.9 | 17.1 ± 3.8 | 3.8 ± 0.4 | 63.7 | 36.3 | 21 | 12 | 201 | 113 |
| Fe-Precise | 10.6 ± 2.4* | 13.5 ± 4.2 | 3.8 ± 0.4** | 57.1 | 42.9 | 6 | 6 | 94 | 63 |
| Cu-Imprecise | 12.1 ± 2.9 | 16.6 ± 3.2 | 3.9 ± 0.4 | 73.3 | 26.7 | 24 | 8 | 136 | 75 |
| Fe-Imprecise | 7.3 ± 2.2* | 11.2 ± 3.8 | 3.1 ± 0.4** | 71.6 | 28.4 | 22 | 13 | 184 | 116 |

\**P*-value for difference in means = 0.034
\*\**P*-value for difference in means = 0.062

### Receivers of signal (dance followers)

Followers tended to exit the nest immediately after following a dance that was relatively precise, but as precision decreased, followers tended to switch to another dancer rather than exit the nest (Fig 3, Table 1) (*z* = −2.24, *P* = 0.025). For each 1° increase in SD of the direction component of a dance, the odds of a follower exiting the nest (versus odds of switching dances) decrease by 6% (95% CI: 0.7% to 10.1%; Fig 3b). The probability a follower would exit the nest increased as the number of waggle phases per dance increased (interaction term: *z* = 2.71, *P* = 0.007; Fig 4), but only for those following sleep-restricted dancers (steel-tagged: *z* = 3.824, *P* = 0.00013; copper-tagged: *z* = 0.573, *P* = 0.57). Followers appeared to follow more waggle phases in a dance if the dance was precise, but, again, only for those following sleep-restricted dancers (Fe-precise vs. Fe-imprecise; marginally nonsignificant at *P* = 0.062; Table 1). The number of followers did not change in response to dance precision (*z* = −0.702, *P* = 0.62), nor did the number of followers/waggle phase (*z* = −0.335, *P* = 0.88).

**Fig 3.**
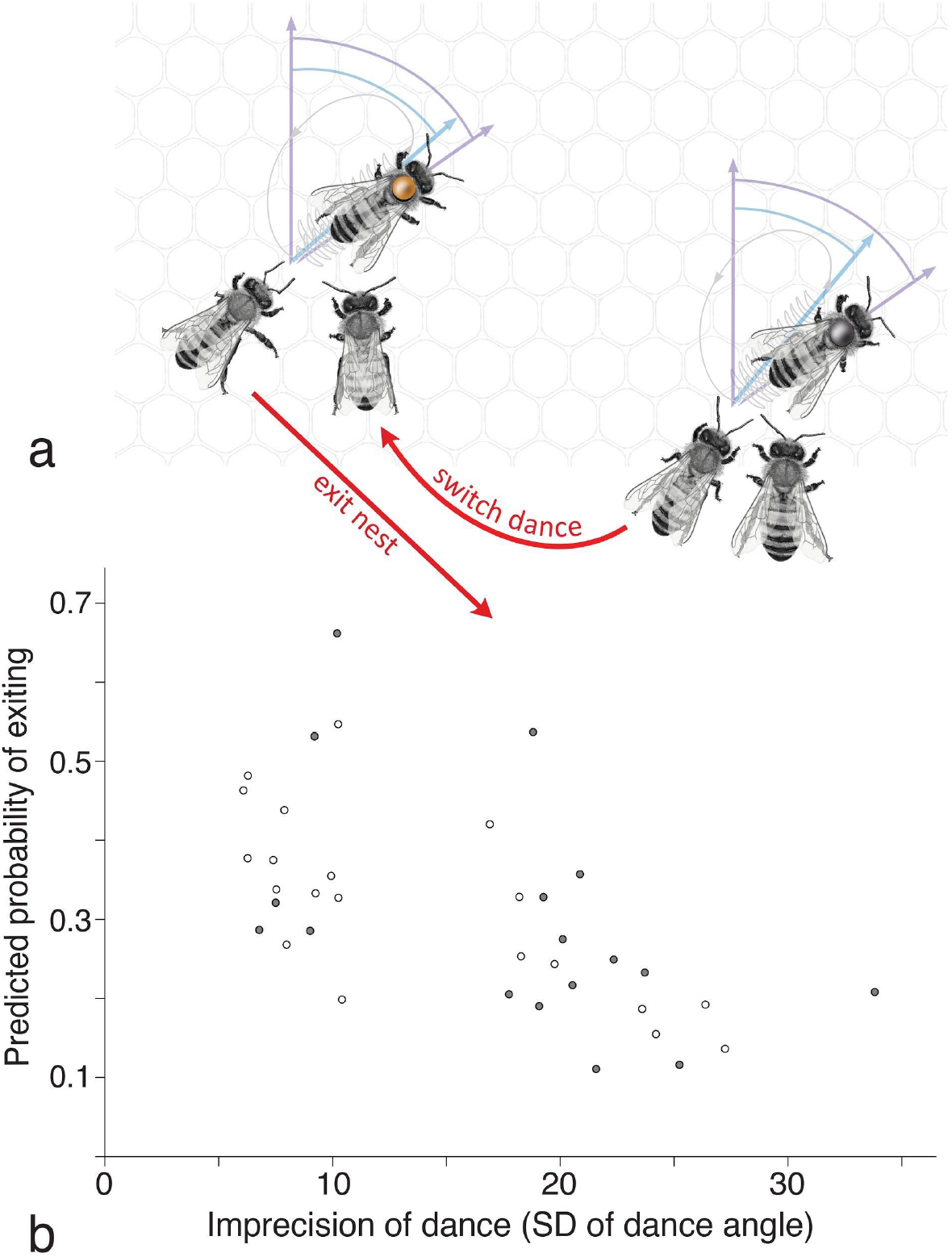
Behaviour by dance followers with respect to dance precision, based on the standard deviation (SD) of the directional component of the dance. (a) Dance followers of a relatively precise dance (left) were more likely to exit the nest than to switch to another dance. Followers of a relatively imprecise dance (right) were less likely to exit the nest and more likely to switch to another dance. (b) Predicted probability of followers exiting the nest based on standard deviation, SD, of the directional component of the dance. The greater the SD, the greater the imprecision. Follower data were collected for dances with SD <10.45° and ≥16.45°, resulting in a gap in the scatter plot (*N* = 39 dances). Gray circles = followers of sleep-restricted, steel-tagged dancers; white circles = followers of control, copper-tagged dancers. Predicted probabilities of switching would be found by subtracting the probabilities of exiting from 1, producing a graph with a trend that is increasing. Honeybee illustrations by Danielle VanBrabant.

**Fig 4.**
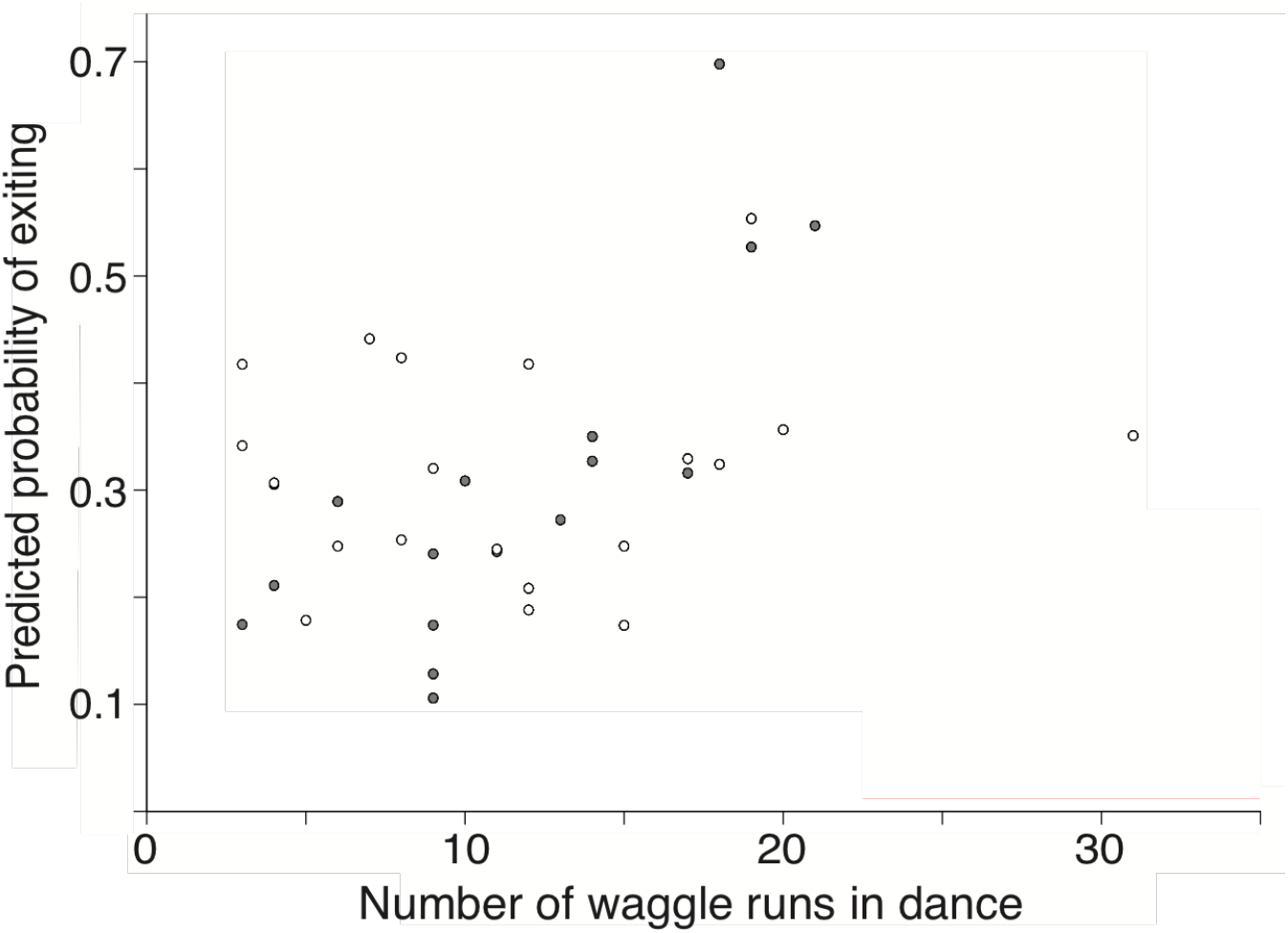
Predicted probability that a dance follower will exit the nest with respect to the number of waggle phases in the dance (*N* = 39 dances). Gray circles = followers of sleep-restricted, steel-tagged dancers; white circles = followers of control, copper-tagged dancers. Predicted probabilities of switching would be found by subtracting the probabilities of exiting from 1, producing a graph with a trend that is decreasing.

The number of waggle phases a bee follows in successive dances could indicate the potential success of switching from one dance to a subsequent dance. In a sample of bees following imprecise dances switching to precise dances, 10 followed more waggle phases and 8 followed fewer. In contrast, of the followers making the opposite switch (precise to imprecise dances), 11 followed more waggle phases and 19 followed fewer. Switching seemed to result in following fewer waggle phases if a follower switched to a less precise dance, or to the dance of a sleep-restricted bee, although neither result is statistically significant.

### Signalers (dancers)

Sleep-restricted dancers not only performed relatively imprecise dances (Fig 5) (SD: 16.2 ± 0.7° vs. 13.6 ± 0.6°; *z* = 2.73, *P* = 0.012; *N* = 50 dances by 6 Fe bees & 76 dances by 7 Cu bees, respectively; linear mixed model with treatment as a fixed effect and dancer as random factor), but may perform fewer dances (7.3 ± 2.1 vs. 12.5 ± 1.4; *t* = 2.07; marginally nonsignificant at *P* = 0.064; *N* = 7 and 6 bees, respectively), and proportionally fewer precise dances than control bees (6 of 28 dances, 21.4%, vs. 21 of 45 dances, 48.9%; *P* = 0.052; Fisher Exact Test; Fig 2, Table 1). Directional precision of dances increased with increasing number of waggle phases, with a marginally nonsignificant increase in dances by control bees (*P* = 0.051), and a stronger relationship in sleep-restricted bees (*P* < 0.0001); for each additional waggle phase by a sleep-restricted dancer, the average SD decreased by 1.52 degrees.

**Fig 5.**
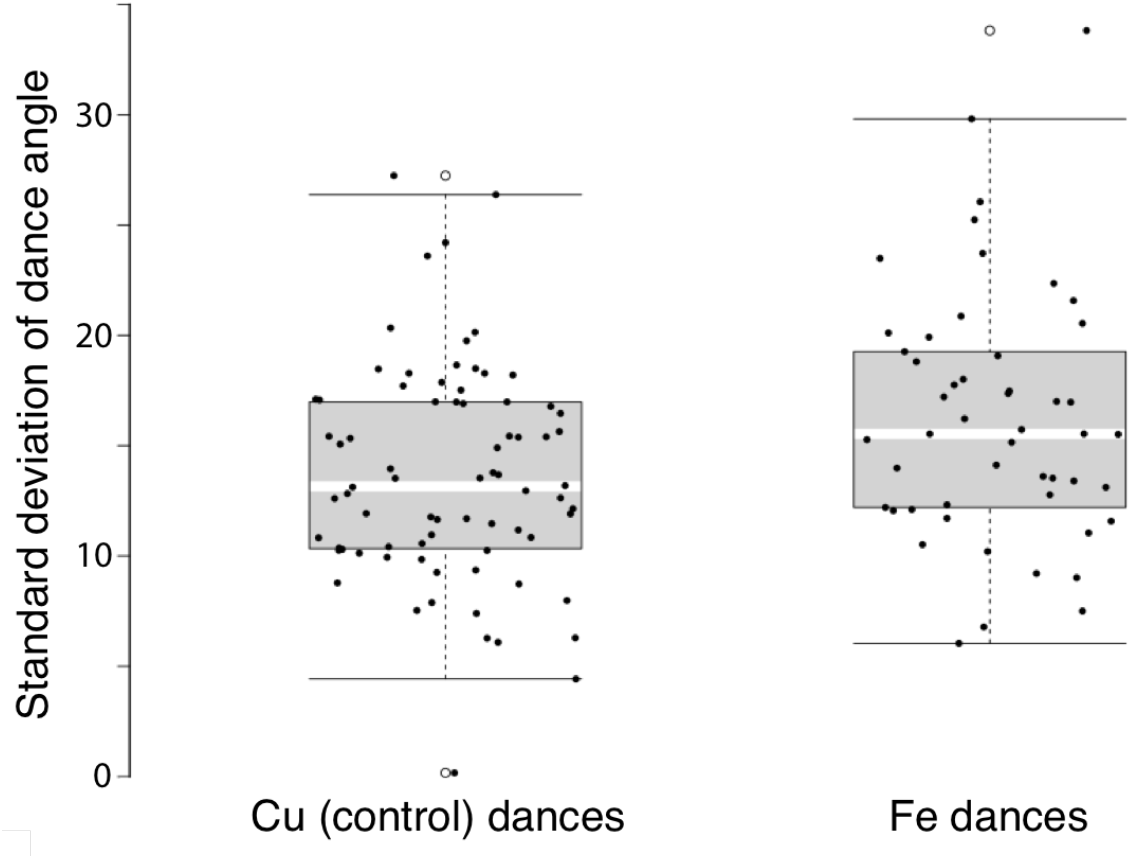
Standard deviation (SD) of dance angles for each dance, by control, copper-tagged dancers, and sleep-restricted, steel-tagged dancers (*N* = 126 dances). These data represent the complete set of dances analysed during the day following sleep restriction, previously displayed as means in Klein et al. (2010; Fig 4).

## Discussion

### Dance precision and follower behaviour

Followers of precise dances were more apt to exit the nest, and followers of imprecise dances were more likely to switch dancers. It might pay for a follower to switch when the perceived quality of the advertised food source is low, perceived risk or cost associated with the journey is high, or degree of variance/error in the signal is too great. In such cases, a follower might have greater success attempting to retrieve food advertised by a different dancer. Because our food source was consistent in quality and location throughout the study, variance in the signal was a more likely factor affecting follower behaviour. If signal variance is an important criterion assessed by a follower, switching to another dancer could increase the probability of obtaining more precise directions, at the expense of time lost that could have been spent foraging.

Multiple bees followed each dance, whether the dance was precise or imprecise. Ultimately, it should not matter how many followers are attracted to a dance, or how many bees follow per waggle phase of a dance, neither of which we found to be related to dance precision. For a dance advertising a food resource to be more successful, it needs to attract more followers that exit the nest and successfully forage, and dance precision was a good indicator of exiting the nest. Exiting the nest did not guarantee that a bee was following the directions of a waggle dance for our feeder, but it is unlikely, for example, that a bee used private information to fly to an alternate food source, considering the dearth of natural food sources available to honeybees at our chosen study site.

There are several factors affecting a dance’s precision, including proximity to the nest. A dance’s directional precision increases predictably as the distance to the advertised source increases from the nest, with the arc representing the foraging area remaining constant (Towne & Gould, 1988; Tanner & Visscher, 2006). Increased precision for more distant sources could be an adaptation, increasing the probability that a follower locates the source (Towne & Gould, 1988; Weidenmüller, & Seeley, 1999; Gardner, Seeley, & Calderone, 2007). Alternatively, the difference in precision could merely be due to physiological constraints of the dance (Tanner & Visscher, 2006; Tanner & Visscher, 2008; Tanner & Visscher, 2010a; Tanner & Visscher, 2010b; Preece, & Beekman, 2014). Precision also differs with respect to what is being advertised – when dancing for a nest site, precision will be greater than when dancing for a food resource (Weidenmüller & Seeley, 1999). Even a sudden increase in food reward (increased sucrose concentration) can affect directional variation, at least at the outset of a dance (De Marco, Gil, & Farina, 2005). None of these factors could have played a role in follower behaviour in our study, because, again, we offered a single resource of consistent quality at a single location.

Since precision is assessed by attending iterations of a dance, how many waggle phases does a bee need to follow to make it to the site being advertised? Followers reduce signal noise by averaging several waggle phases for an overall vector (Tanner & Visscher, 2008), and the number of waggle phases followed correlates positively with accuracy of the flight path (Tanner & Visscher, 2009), so unless the perceived risk involved is greater (Wray, Klein, & Seeley, 2011), following more waggle phases would appear to benefit a follower. Toufailia et al. (2013) showed that workers follow fewer waggle phases as distance to food location increases, but followers invest more time when following a dance for a more distant site because each waggle phase and return phase takes longer. The mean number of waggle phases followed to reach an advertised site ranges widely, with Judd (1995) reporting on the lower end of 7.8 ± 3.2 (SD) consecutive waggle phases followed before making it to a feeder 150 m away, but naïve followers have successfully reached a destination after following only six (200 m away), five (400 m away), or as few as two waggle phases (*N* = 2 followers, 150 m away) (Esch & Bastian, 1970; Mautz, 1971; Judd, 1995; Couvillon et al., 2012). Other cases of successful flights following so few waggle phases can be due to “reactivation,” when followers have experience with the advertised site and respond to the odor of the dancer, rather than the precision of the dance (Wray et al., 2011; Toufailia et al., 2013). From the perspective of a human decoding a dance, Couvillon et al. (2012) concluded that four consecutive waggle phases occurring in the middle of a dance represents closely the mean for all of the waggle phases within the dance. We found that our bees followed approximately this number of waggle phases (Table 1) for our feeder, located 1 km from the nest. It is important to note that foragers follow several dances (4.8 ± 3.2; range: 1-12) and make several excursions in search of the advertised flower patches before successfully locating a natural recruitment target (Seeley, 1983). Precision may make a difference with regard to not only how many waggle phases are necessary to follow, but how many dances are followed before recruitment is successful.

Honeybee workers can be observed to follow several different dances before exiting the nest, though one study indicates they follow only one dance closely (Seeley & Towne, 1992). Whether or not followers compare dances, if they reject some by switching, and exit after following others, dance success hinges on the factors guiding the decision to exit or switch. We found that dance directional precision relates to the outcome of a follower’s decision to exit or switch, as does number of waggle phases per dance, but other dance variables appear to affect follower behaviour as well (Grüter & Farina, 2009). These variables include substrate vibration (Tautz, 1996; Tautz & Rohrseitz, 1998), airborne sound (Michelsen, Andersen, Kirchner, & Lindauer, 1989; Kirchner & Sommer, 1992), tactile cues (Rohrseitz & Tautz, 1999), semiochemicals (Thom, Gilley, Hooper, & Esch, 2007), or a dancer’s excitement, expressed as thoracic vibrations (Hrncir, Maia-Silva, Cabe, & Farina, 2011). Food quality can affect duration of the return phase of a waggle circuit and the number of waggle phases per dance (Seeley, Mikheyev, & Pagano, 2000), which also relates to the outcome of a follower’s decision. However, with no variation in food source, we found no difference in duration of return phase between relatively precise and imprecise dances (3.4 ± 1.9s vs. 6.4 ± 8.4s SD), using durations starting with the second return phase (max 5 phases total, if dance consisted of <7 phases). Changes in exuberance may be a worthwhile place to look for effects of sleep restriction on dancer behaviour and consequential behaviour of the dancer’s followers.

Bees occasionally dance with errors of 10-15° from the solar angle, yet followers make it to their destinations (von Frisch, 1967). The mean error within our sample of imprecise dances was 21.1° (*N* = 46 dances), which translates into an error of 366 m from the food source (law of cosines, given that the food source was 1 km from the nest) for one SD of angular variability. Dancers tend to alternate between waggle circuits that begin with left and right turns, and sleep-restricted dancers’ precision decreased during waggle phases specifically after left turns (Klein et al., 2010), suggesting that direction would be averaged as an inaccurate vector across all (left and right) waggle phases. Imprecision could be problematic for a follower if the averaged angle results in an inaccurate vector. Flight efficiency depends on managing energy costs and gains (Stabentheiner & Kovac, 2016), so additional flight time with reduced probability of arriving at a food source could make imprecise dancing a detriment. Metabolic costs associated with additional flight time can be converted into calories (Seeley, 1983), shedding estimable light on the cost of following imprecise dances. If costs of searching are considerable, benefits of receiving accurate and precise information from the dancer are probably considerable as well. What a bee experiences outside the nest, including floral odors (von Frisch, 1967), and the social behaviour of experienced foragers (Grüter & Farina, 2009) likely reduces recruits’ flight time by helping to pinpoint the food source. To what extent these mitigating factors compensate for imprecision or inaccuracy is unknown.

Followers of an imprecise dance tended to switch dances, offering them opportunities to obtain more precise directions, especially if they had followed fewer waggle phases previously. Following fewer waggle phases gives a less complete, and less accurate picture of an advertised location (Tanner & Visscher, 2009), and fewer waggle phases were followed in the relatively imprecise dances performed by our sleep-restricted dancers. Although switching dances offers the possibility of obtaining increased directional precision, sleep-restricted dancers perform potentially fewer dances, proportionally fewer precise dances, and these dances include fewer waggle phases per dance, so the opportunities for success at the colony level decrease dramatically, particularly if within this reduced rate of dancing, followers are switching dances. Switching dances means postponing or forfeiting departure from the nest and, ultimately, possibly reducing efficiency of food acquisition. Lost efficiency due to corrupted communication could render a colony less fit than colonies that experience a sounder night’s sleep. As we discovered, the follower behaviour could not be explained entirely by imprecision caused by dancer sleep loss, however. When a dancer had experienced sleep loss, follower switching behaviour was best explained by total number of waggle phases in the dance, and number of waggle phases followed was best explained by dance precision. Because switching by followers did not significantly correlate directly with dance precision based on a dancer’s sleep loss, followers must have responded to cues associated with sleep-restricted dancers apart from the directional precision of their dances. The honeybee waggle dance is one of the best studied and most celebrated examples of animal communication, and sleep-related effects on dance communication could lend insight into connections between sleep and communication in other signaler-receiver, or information-processing scenarios.\

---

**Movie S1.**
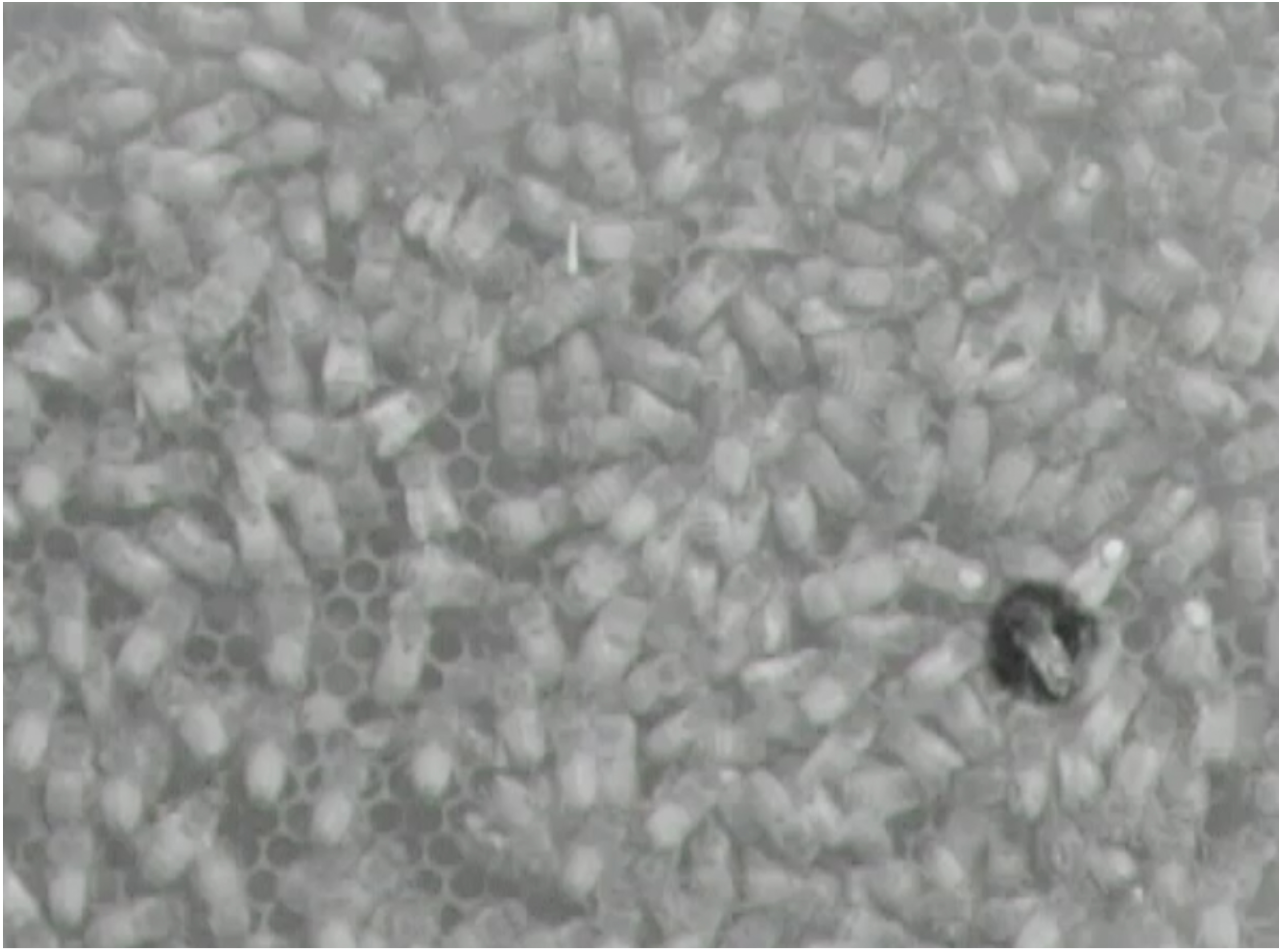
Dance followers, exiting the nest and switching to another dance. Tunnel leading to hive exit is located immediately off-screen at lower right. Clip 1: Dance follower exits nest after following precise waggle dance, performed by copper-tagged, control dancer. Follower (#10, highlighted using Adobe After Effects CS6, v. 11.0.4.2) followed dancer (gb, dance #177, with 19 total waggle phases) for 8 waggle phases, lasting 24 s, before exiting the nest. Precision of dance angle for dance followed: SD = 10.25° (<10.45° was considered precise). Clip 2: Dance follower switches dances after following imprecise waggle dance, performed by a sleep-restricted dancer. Follower (#11, highlighted using Adobe After Effects CS6, v. 11.0.4.2) followed dancer (ry, dance #201, with 9 total waggle phases) for 3 waggle phases, lasting 12 s, before following another dancer for 3 waggle phases. Precision of dance angle for first dance followed: SD = 23.71° (≥16.45° was considered imprecise).

**Dataset S1.** Complete followers: all data organized by followers of waggle dances [*N* = 332 follower events, including 169 follower events by 5 Fe bees in 14 dances (5 precise dances, 9 imprecise dances) and 163 follower events by 5 Cu bees in 13 dances (8 precise dances, 5 imprecise dances)]. Original file name, used in R code: FollowersPlusReduced

**Dataset S2.** Followers of sleep-restricted dancers: data organized by followers of sleep-restricted, Fe waggle dancers [subset of S1 Data; *N* = 169 follower events by 5 Fe bees in 14 dances (5 precise dances, 9 imprecise dances)]. Original file name, used in R code: FollowersPlusReducedFe

**Dataset S3.** Followers of control dancers: data organized by followers of Cu waggle dancers [subset of S1 Data; *N* = 163 follower events by 5 Fe bees in 13 dances (8 precise dances, 5 imprecise dances)]. Original file name, used in R code: FollowersPlusReducedCu

**Dataset S4.** Frequency of follower events [*N* = 37 dances, including 17 dances by 5 Fe bees (5 precise dances, 12 imprecise dances) and 20 dances by 6 Cu bees (12 precise dances, 8 imprecise dances)]. Two dancers were “lost” during video observations. We excluded these dancers and their two dances from the analysis since these data didn’t provide any information about exiting of switching, hence the difference between sample size here and that mentioned in Results. Original file name, used in R code: followerfreqPlus

**Dataset S5.** Complete waggle dance data (*N* = 126 dances, including 51 dances by 7 Fe bees and 75 dances by 6 Cu bees). Original file name, used in R code: allDancesPost

**Dataset S6.** Waggle dance data, used to test the average number of dances per dancer (7 Fe bees, 6 Cu bees). Original file name, used in R code: allDancesNumberperDancer

**Dataset S7.** Precise waggle dance data: dances with a relatively precise directional component (6 dances by Fe bees and 21 dances by Cu bees). Original file name, used in R code: allDancesPostBinnedP

**Dataset S8.** Imprecise waggle dance data: dances with a relatively imprecise directional component (22 dances by Fe bees and 24 dances by Cu bees). Original file name, used in R code: allDancesPostBinnedI

**Code S1.** R code used for analyses of dances and dance follower behaviour.

